# Vulnerability to cavitation is linked to home climate precipitation across eight eucalypt species

**DOI:** 10.1101/2021.09.05.459049

**Authors:** David Coleman, Andrew Merchant, William T. Salter

## Abstract

Vulnerability to cavitation in leaves is the result of highly adaptive anatomical and physiological traits that can be linked to water availability in a species’ climate of origin. Despite similar gross leaf morphology, eucalypt species are often confined to specific climate envelopes across the variable rainfall environments of Australia. In this study, we investigate how the progression of cavitation differs among eucalypts and whether this is related to other hydraulic and physical leaf traits. We used the Optical Visualisation technique to capture cavitation progression across the leaves of eight eucalypt species (*Angophora crassifolia, Corymbia tessellaris, Eucalyptus atrata, Eucalyptus grandis, Eucalyptus laevopinea, Eucalyptus longifolia, Eucalyptus macrandra, Eucalyptus tereticornis*) from a wide range of climates and grown in a common garden setting. Vulnerability to cavitation, represented by the leaf water potential required for 50% cavitation of leaf vessels, varied significantly among species (−3.48 MPa to −8.25 MPa) and correlated linearly with home climate precipitation and leaf SLA (*R*^2^ of 0.64 and 0.75, respectively). P12-P88, the range of water potentials between which 12% to 88% of cavitation occurs, was decoupled from P50 but also correlated with leaf SLA (*R*^2^ of 0.72). We suggest the magnitude of P12-P88 may be representative of a species’ drought strategy – a large P12-P88 signifying leaves that exhibit drought tolerance (retention of leaves under drought conditions) and a small P12-P88 signifying drought avoidance (leaf shedding after a threshold of drought is reached). Our results agree with other studies that highlight these cavitation metrics as genetically fixed traits. Turgor loss point, on the other hand, may be more plastic, as evidenced by the low variability of this trait across these eucalypt species grown in a common garden environment. Further study will help to establish the SLA-related anatomical traits that impart cavitation resistance and to extend these conclusions to a greater number of species and home climates.

## Introduction

A primary driver of evolutionary adaption in land plants is water availability in the environment, leading to a wide diversity in hydraulic structure and function across plant species. Diversity is most apparent in leaves, which must maximise surface area to intercept light while minimising water loss to transpiration. As the site of evaporation between the plant and the atmosphere, leaves of land plants must also contend with more negative water potentials (Ψ_leaf_) at earlier stages in plant hydraulic stress than other organs (Scoffoni *et al*., 2018). If adequate water is not available to replace that lost to transpiration, leaf tissues face damage due to desiccation and loss of function. Cavitation or embolism, a phase change from liquid water to gas within xylem conduits under high water tension, is the dominant mechanism by which desiccation-induced tissue damage spreads through a plant (Choat *et al*., 2018). In leaves, this causes an interruption in the supply of water to the mesophyll (Sack and Holbrook, 2006). Although embolism repair is possible (Klein *et al*., 2018), loss of hydraulic integrity in leaves is often associated with delayed or non-reversible damage following periods of hydraulic stress (Skelton *et al*., 2017). The ability of plants to resist the process of cavitation and maintain hydraulic integrity in the leaf vein network dictates the survival of the leaf tissue and, ultimately, the productivity of the plant. Therefore, cavitation resistance of leaves under hydraulic stress is a key trait with which species can be compared (Brodribb *et al*., 2016a).

There are general rules that apply to the behaviour of angiosperm vein networks under hydraulic stress. Conduits in large veins are the first to cavitate and appear to do so multiple times, suggesting redundancy in the hydraulic pathways connecting sections of the mesophyll (Brodribb *et al*., 2016a). Redundancy in leaf vein networks imparts resilience to the system by providing alternate routes for water to reach distal parts of the mesophyll during hydraulic stress and acts as an insurance policy to prevent rapid and catastrophic failure of tissue structure. Therefore, gradual cavitation over larger ranges of Ψ_leaf_ suggests greater tolerance to hydraulic stress (Brodribb *et al*., 2016a).

A tradeoff between efficiency and strength has been found to exist in the construction of plant xylem networks in leaves (Liu *et al*., 2021; Grossiord *et al*., 2020; Pritzkow *et al*., 2019). Thinner, more numerous conduits have increased tolerance to cavitation than larger conduits but have a higher mass to flow area ratio, increasing the hydraulic resistance of flow through the leaf and costing the plant more in the tradeoff to carbon fixation and evaporative cooling (Santiago *et al*., 2018; Klein *et al*., 2018). In addition to vein traits, many other anatomical and physiological leaf traits are associated with cavitation resistance. Thicker leaves provide a buffer to rising tension inside xylem conduits with declining leaf water status, with larger outside-xylem water stores protecting against embolism (Scoffoni *et al*., 2017a). Drought resistance traits, such as low turgor loss point (TLP), often cooccur with resistance to cavitation in leaf vein networks. The onset of leaf hydraulic dysfunction also corresponds with full stomatal closure in plants exposed to gradual drought treatment in some species (Johnson *et al*., 2018; Li *et al*., 2019). In scenarios of rapid leaf desiccation (hours rather than days or weeks), stomatal closure is thought to occur well before the leaf suffers significant damage (Li *et al*., 2019; Trueba *et al*., 2019). However, Skelton *et al*. (2015) reported that some plants have small or negative differences between Ψ_leaf_ at stomatal closure and the leaf xylem cavitation threshold, the stomatal hydraulic safety margin.

In addition to leaf-level traits and physiological processes that influence cavitation resistance, leaves also operate in concert with the hydraulic status of the whole plant. In order to draw ecophysiological conclusions from cavitation resistance in leaves, it is essential to acknowledge differing plant survival strategies in the face of drought stress, such as the hydraulic vulnerability segmentation hypothesis (Tyree & Ewers, 1991). In many cases, xylem vessels in leaves have been shown to be more vulnerable to hydraulic decline than stems, evidenced by the high ratio of the water potential at 50% stem xylem cavitation (P50_stem_) to leaf xylem cavitation (P50_leaf_), the safety margin of hydraulic segmentation (Pivovaroff *et al*., 2014a). Where this is the case, leaves can be seen to act as comparatively expendable “hydraulic fuses” during drought, acting to delay or prevent loss of sapwood conductivity as this is more critical to the survival of the plant. Interpreting P50_leaf_ in these species to be solely representative of the vulnerability of the whole plant to drought stress would not be accurate. Although this hydraulic segmentation hypothesis certainly applies to the leaves of some species, there appears to be a range of different degrees to which P50_leaf_ differs from P50_stem_ that correlate with climate and biome metrics, including many plants that possess negligible or even negative ratios in wetter, more humid environments (Zhu *et al*., 2016). At the very least, leaf hydraulic vulnerability is critical to plant survival when we consider that leaf cavitation will lead to tissue death, which ultimately limits productivity (Pritzkow *et al*., 2019).

Cavitation across leaf venation networks can be visualised and quantified using several techniques, including the use of cryo-scanning electron microscopy (Johnson *et al*., 2009) and micro-CT reconstructions (Scoffoni *et al*., 2017b) to map the formation of air spaces inside vessels (Scoffoni *et al*., 2017b). However, these imaging techniques rely on expensive equipment and specialist expertise. Optical visualisation (OV), on the other hand, is a low-cost and easy-to-use alternative in which cavitation events can be detected from a sequence of standard photographic images of a drying leaf. Rapid, embolism-induced changes are distinguished via a difference in the transmission of light through the tissue in successive images (Brodribb *et al*., 2016b). Coupling the OV technique with a measure of leaf hydraulic status allows for the construction of vulnerability curves, which describes a species’ resistance to embolism. Hydraulic vulnerability curves show the percentage loss of conductivity (PLC) as a function of decreasing xylem pressure and can be used to extract metrics of vulnerability – various Ψ_leaf_ thresholds beyond which significant xylem embolism occurs (Venturas *et al*., 2017). The most commonly used parameter, P50 (the bulk leaf water potential at which 50% loss of conductivity has occurred), can be used to summarise the complex progression of cavitation propagation, with more negative P50 indicating vein networks more resilient to hydraulic stress (Blackman *et al*., 2017; Hartmann *et al*., 2018). However, the characteristic sigmoidal shape of hydraulic vulnerability curves (Creek *et al*., 2019) means other metrics such as P12 (a measure of the initiation of significant xylem embolism) and P88 (completion of xylem embolism) are also helpful to compare strategies to hydraulic stress across species. For example, species that initiate cavitation earlier in a desiccation treatment than other species (higher P12) but fully cavitate later (lower P88) suggests a different drought strategy to leaves in which cavitation occurs over a narrow range of water potentials. We hypothesize that the P12-P88 range could be used as a new metric to define whether species tolerate drought (long and gradual cavitation progression starting at low water stress) or attempt to avoid it (rapid and complete cavitation progression at a particular threshold water stress).

Eucalypts are a group of woody angiosperms ideally suited to drawing ecophysiological links between the environment and leaf hydraulic traits (Merchant *et al*., 2007). The term eucalypt refers to a grouping of species across three closely-related genera of the Myrtaceae family – *Eucalyptus, Corymbia* and *Angophora*. Across Australia, eucalypt species have adapted to a wide range of climates with varying levels of water availability while maintaining relatively similar gross leaf size and shape (Brooker and Nicolle, 2013). In fact, similar and flexible gross leaf morphology within and between species has prompted the scientific community to use microscopic venation patterns as a tool for differentiating species in these genera (Brooker and Nicolle, 2013). Therefore, it is possible that the pressure thresholds that hydraulic structures in leaves can withstand may contribute to confining species to a particular climate envelope (Blackman *et al*., 2012). Recent research has related the hydraulic vulnerability of multiple eucalypt species to their home climate (Li *et al*., 2019; Bourne *et al*., 2017), including the safety margin of hydraulic segmentation (water potential at stomatal closure to P12) and the Hydroscape area (the region bounded by the predawn water potential (Ψ_pd_) and midday water potential (Ψ_md_) regression and 1:1 line on a Ψ_pd_ vs Ψ_md_ plot) (Li *et al*., 2019). However, a detailed study of the spatial progression of leaf embolism and how this relates both to physical leaf properties and to home climate across eucalypt species could reveal new insights into the drivers and strategies employed to mitigate desiccation.

In this study, we measured the progression of cavitation in response to declining water status in leaves of eight eucalypt species from home climates spanning an extensive geographic range and different subgenera, grown in a common garden setting. We used the OV method (Brodribb *et al*., 2016a) to observe the spatial and temporal dynamics of hydraulic vulnerability across the leaf. We quantified key cavitation metrics (including P50 and TLP) and measured other important hydraulic and physical leaf properties to determine the adaptive mechanisms underpinning leaf cavitation resistance across these eight species. Specifically, we aimed to investigate (i) whether the characteristics of embolism progression are consistent across these eucalypt species, (ii) whether cavitation metrics correlate with other leaf thresholds of hydraulic decline, such as TLP, and (iii) which leaf traits or home climate variables confer cavitation resistance across eucalypt species.

## Methods

### Site characteristics

Eight species of eucalypt were chosen for study encompassing a broad taxonomic scope, including representatives from the *Angophora, Corymbia* and *Eucalyptus* genera – *Angophora crassifolia, Corymbia tessellaris, Eucalyptus atrata, Eucalyptus grandis, Eucalyptus laevopinea, Eucalyptus longifolia, Eucalyptus macrandra* and *Eucalyptus tereticornis*.These species originated from a wide range of climates (Table 1). Of note is the range of mean annual temperature (13.2 °C for *E. laevopinea* to 23.5 °C for *C. tessellaris*) and mean annual precipitation (503 mm for *E. macrandra* to 1335 mm for *E. grandis*) of seed collection locations.

**Table 1.**
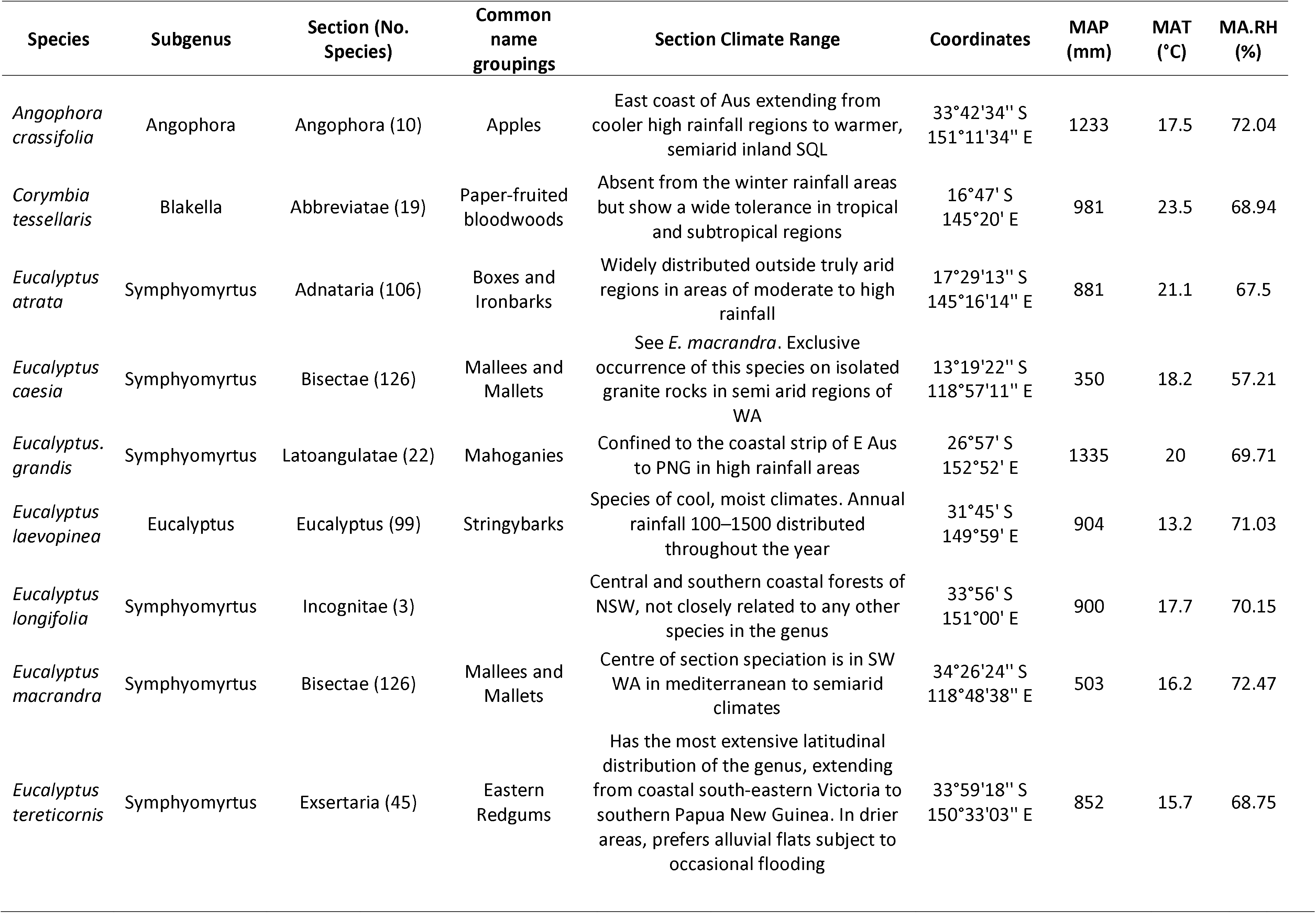
Summary of species taxonomy, short description of the section distribution, location of seed collection locations, mean annual temperature (MAT), mean annual precipitation (MAP) and mean annual relative humidity (MA.RH) of seed collection locations.

All plant material was sourced between mid-June to mid-August 2020 from the Australian Botanic Garden, Mount Annan, NSW, Australia (34° 4′13″S, 150°46′12″E, elevation 150m) (See Supp. Figure 1 for map location). The site has a mean annual temperature of 16.5°C and mean annual precipitation of 780 mm. During the six months prior to the experiment, temperatures were close to the long-term average for this time of year (average daily maximum = 20-30°C, average daily minimum = 10-20°C) but rainfall was almost double the long-term average (pre-experiment = 566 mm, long-term average = 295 mm). Temperatures and rainfall during the experiment were similar to the long-term average, with daily average maximum and minimum temperatures of 17-18°C and 3-5°C, respectively, and ^~^100 mm rainfall over the 2.5 months. The soils in the Garden are dominated by well-drained Kurosols and Dermosols. Seed progeny data was provided by The Australian Botanic Garden (Table 1) (See Supp. Figure 1 for map location). Home climate data for species were sourced from the Atlas of Living Australia Spatial Portal (http://spatial.ala.org.au/; accessed 6 October 2020).

### Sample collection

Each morning of the experiment, one small branch (^~^ 30 cm) of adult foliage was cut from a tree using clean sanitised secateurs. Sampled branches were always situated on the light exposed, north or east-facing side of the tree. The branch was immediately recut underwater, covered, and transported (20 minutes) by car to a laboratory at the University of Sydney, Brownlow Hill, NSW. The branch was then placed in a dark cupboard to rehydrate, and leaf surfaces dried with a tissue, if necessary, until measurement (1.5 – 2 hours after initial cutting).

### Measurement of cavitation, water potential and hydraulic vulnerability metrics

Progression of leaf vein cavitation (leaf hydraulic vulnerability) with declining Ψ_leaf_ was measured using the open-source optical visualisation technique (Brodribb *et al*., 2016b). In the laboratory and over the course of the experiment, three to four healthy fully expanded leaves from different individual trees of each species were positioned and held in place by custom-built leaf clamps underneath a digital camera (Camera Module v2; Raspberry Pi Foundation, Cambridge, UK) and macro-lens (Eschenbach 20x Magnifier; Prospectors, Sydney, Australia). Refer to www.opensourceov.org for a comprehensive description of the design. Leaf edges were taped to the surface of the clamp to avoid movement and shrinkage during the experiment. The water source to the branch was then removed and the leaf, still attached to the branch, was left to dry on the lab bench. Images of the leaf within the clamp were taken at 1 min intervals and stored on a Raspberry Pi computer (3B+; Raspberry Pi Foundation, Cambridge, UK). Leaf water potential (Ψ_leaf_) was monitored continuously during leaf desiccation using a psychrometer (PSY1 Stem Psychrometer, ICT International, Armidale, NSW, Australia) at 10 minute intervals. The psychrometer was installed on an adjacent leaf rather than the stem, due to the small stem width of the branchlet. The psychrometer was installed in a similar way as would be used on the stem, abrading a small area of the leaf cuticle, clamping the psychrometer to the tissue, sealing the instrument against the surface of the leaf using vacuum grease and insulating the apparatus using packing foam and a cloth (Dixon and Downey, 2013). The temperature in the lab remained between 20 and 23 degrees over the course of the experiment and temperature changes recorded by the psychrometer were within 0.1 degree between successive measurements recorded at 10 minute intervals.

Leaf image sequences were analysed using an image subtraction method in ImageJ (Rueden *et al*., 2017) with the Fiji distribution (Schindelin *et al*., 2012). To visualise and quantify cavitation, we used the OSOV Toolbox and image processing methodology (Brodribb *et al*., 2016b), available at www.opensourceov.org. Approximately 2000 captured colour images for each replicate were converted to black and white 8-bit images. Pixel values of consecutive images were subtracted from each other to reveal changes using the “Image difference” function of the OSOV add in. The “moments” thresholding mask and the “despeckle” image processing functions were used to reduce background noise and noise from leaf shrinkage. Given that this noise varied from leaf to leaf, these functions were applied manually such that only the pixels representing cavitating leaf veins were highlighted. The total number of pixels highlighted on each image was quantified using the “measure stack” function and the percentage cavitation over time given as the ratio of the cumulative pixel sum to the total pixels highlighted after cavitation had finished.

Hydraulic vulnerability was inferred by combining the Ψ_leaf_ as measured by the psychrometer with the OV data (see Data processing section). Ψ_leaf_ change with time was initially rapid but slowed and became linear following stomatal closure (Brodribb *et al*., 2016b). Despite the linear nature of Ψ_leaf_ decline, a smoothing spline was fitted over the cavitation - Ψ_leaf_ data to match timestamps between the psychrometer and the OSOV images using the *smooth.spline(*) function from the stats package in base R (R Core Team, 2019). A technical limitation of these stem psychrometers was that the lowest measurable Ψ_leaf_ was −7 MPa. However, at these levels of water stress, turgor pressure was already lost, and we assumed the rate of Ψ_leaf_ change would not deviate after this point. Therefore, following Brodribb *et al*. (2016a) and Rodriguez-Dominguez *et al*. (2018), linear regressions were fitted to extrapolate the hydraulic vulnerability of leaves beyond −7 MPa. P50 was defined as the Ψ_leaf_ where 50% of the pixels in the OV images were highlighted. P12 and P88 were calculated in a similar way, at 12% and 88% cavitation area, respectively.

### Pressure-volume (PV) curves

One leaf per branch from each of the same individuals was harvested, hydrated in the dark in the same way as the whole branches and used to construct pressure-volume relationships using a Scholander pressure chamber (Model 1505D, PMS Instrument Company, Albany, USA) and electronic balance to 5 decimal places. Leaf pressure volume (PV) curves were generated following Tyree and Hammel (1972). The leaves were excised, immediately weighed and then periodically weighed and the corresponding Ψ_leaf_ measured as the leaf dried under laboratory conditions. A single PV curve from the four leaves was generated per species for estimation of the turgor loss point (TLP), the bulk elastic modulus (ε) and the osmotic potential at full turgor (Ψ_FT_). TLP was defined as the water potential at the inflexion point of the inverse Ψ_leaf_ – RWC relationship and ε defined as the pre-TLP gradient of the Ψ_leaf_ – RWC relationship.

### Specific leaf area

Five comparable leaves from the same branch were selected based on similar appearance to the measurement leaf and photographed on a flat surface for leaf area calculation using the EBImage package (Pau *et al*., 2010) in Rstudio (Team RStudio, 2015). These leaves were immediately weighed and stored in a 60 °C oven overnight and weighed again to calculate an average specific leaf area (SLA) for the branch using an electronic balance correct to five decimal places.

### Data processing and statistical analyses

All data processing and analyses were conducted in Rstudio (Team RStudio, 2015). The mean and standard error were calculated using Rstatix package (Kassambara, 2020) get_summary_stats() function and significantly different groupings were made using the LSD.test() function in the agricolae package (de Mendiburu and de Mendiburu, 2020). Relationships between climatic and physiological variables were analysed with linear regression using the ggpmisc package (Aphalo, 2016) and the stat_poly_eq() function. Data were visualised using ggplot2 (Villanueva and Chen, 2019).

## Results

### Cavitation progression

Cavitation events in leaf vein networks were visualised for all eight species using the OV method and there was a common pattern in the progression of cavitation among all leaves. Vein cavitation proceeded in large steps initially due to multiple failures of midvein and 1^st^ order vein conduits, followed by smaller, more gradual cavitation of disparate vein networks across the mesophyll (Figure 1). For example, *E. atrata* midvein experienced cavitation at Ψ_leaf_ of −2.5 MPa (blue/purple colour in Figure 1) while the smaller veins of this species cavitate at Ψ_leaf_ of −6 MPa (yellow in Figure 1). Cavitation in multiple conduits of the midvein occurred with increasingly negative water potential in the lateral margins of the midvein in all species and this was particularly evident in *E. grandis* and *E. atrata* (Figure 1). The exception was *E. macrandra* where no midvein cavitation was captured by the OV method. There were visible differences across species in the density of venation structure. Qualitatively, *C. tessellaris* showed cavitation events across a dense, highly reticulated network, while *E. laevopinea* and *E. longifolia* leaves showed fewer cavitation events in lower order veins.

**Figure 1.**
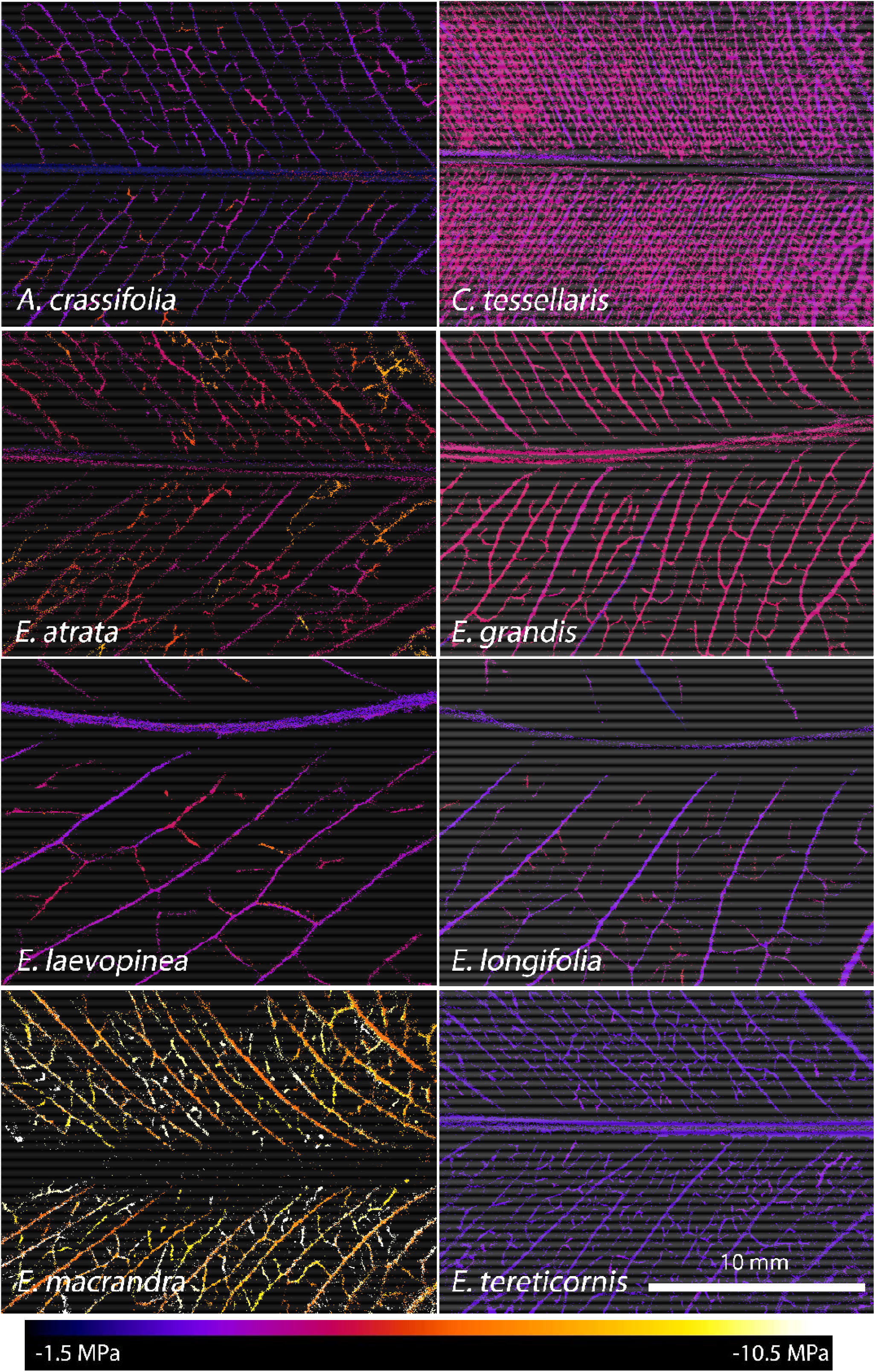
Colour coded cavitation events per 0.1 MPa of leaf water potential for eight species of eucalypt. Colours represent a continuous scale from −1.5 to −10.5 MPa bulk leaf water potential. Images are one representative leaf from each species.

The shape of the hydraulic vulnerability curves within replicates of the same species were reasonably similar (Figure 2). Species that seemed to cavitate over a smaller period of water potentials, such as *C. tessellaris* and *E. grandis*, were particularly tight in their agreement and possessed a sigmoid shape with a steep gradient at the level of P50. Larger differences between individuals were observed in species that exhibited cavitation more gradually, increasing variability in the P50 statistic, such as for *E. laevopinea* and *A crassifolia*.

**Figure 2.**
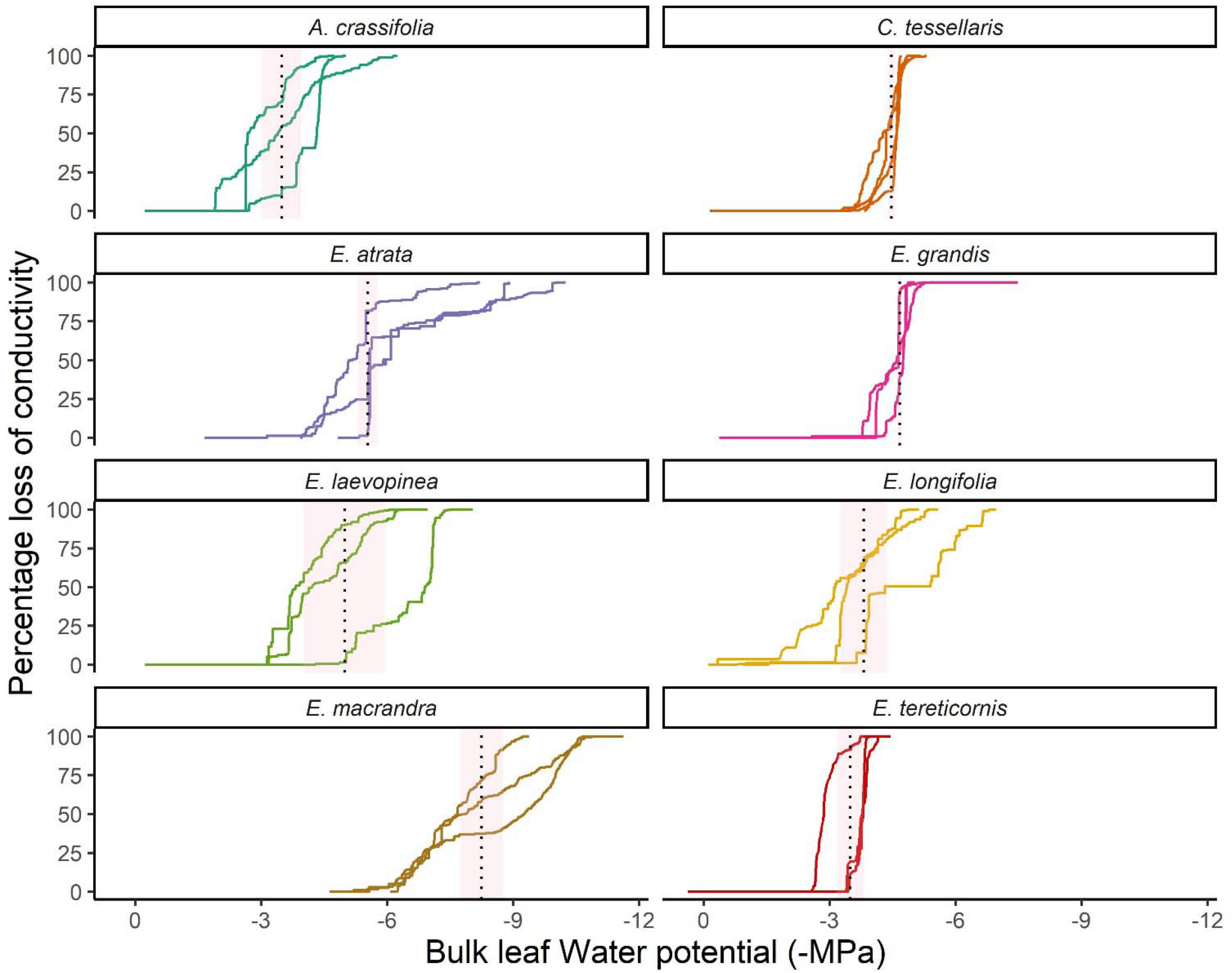
Leaf hydraulic vulnerability curves for eight species of eucalypt. Coloured lines represent the cumulative percentage loss of leaf xylem conductivity due to cavitation over increasingly negative bulk leaf water potential. Dotted lines represent the mean water potential at which 50% of visible cavitation occurred (P50), shaded areas indicate the se. n = 4 for A. crassifolia and C. tessellaris, n = 3 for other species.

### Variation in hydraulic vulnerability between species

The embolism period, designated by the P12-P88 parameter, was found to vary significantly across species (P < 0.05) (Table 2). In some species P12-P88 spanned a narrow range of Ψ_leaf_, such as *C. tessellaris* and *E. grandis* in which P12-P88 were 0.64 ± 0.13 MPa and 0.7 ± 0.2 MPa, respectively. In contrast, this period spanned more than four times that range for other species, for *E. atrata* and *E. macrandra* P12-P88 spanned 2.9 ± 0.78 MPa and 3.19 ± 0.49 MPa, respectively. The P50 threshold was also reached at markedly different Ψ_leaf_ across species (P < 0.05), ranging from −3.48 ± 0.47 MPa in *A. crassifolia* to −8.25 ± 0.51 MPa in *E. macrandra*. Other thresholds of leaf desiccation spanned much smaller ranges of Ψ_leaf_ with little variation across species (P > 0.05), with Ψ_TLP_ spanning a range of 1.1 MPa across all studied species and Ψ_FT_ varying by 1.29 MPa across species. The bulk elastic modulus of leaf tissues (ε) was lowest for *E. laevopinea* (24.39 MPa) and highest for *E. tereticornis* (43.41 MPa) leaves.

**Table 2.** Physiological characteristics and hydraulic vulnerability characteristics for eight eucalypt species. Error ranges are the standard error and groupings are LSD test where significant differences existed between species. MAP: mean annual precipitation, MAT: mean annual temperature, MA.RH: mean annual relative humidity at 3 pm, SLA: specific leaf area, Ψ_TLP_: water potential at turgor loss, Ψ_FT_: water potential at full turgor, ε: bulk elastic modulus, P50: water potential at 50% cavitation, P12-P88: range of water potentials at which cavitation occurred.

| Species | SLA<br>(m<sup>2</sup> kg<sup>-1</sup>) | Ψ<sub>TLP</sub><br>(Mpa) | Ψ<sub>FT</sub><br>(Mpa) | ε<br>(Mpa) | P50<br>(Mpa) | P12-P88<br>(Mpa) |
| --- | --- | --- | --- | --- | --- | --- |
| *A. crassifolia* | 6.05 ± 0.41 ab | -2.74 | -2.65 | 25.03 | -3.48 ± 0.47 a | 1.86 ± 0.57 bc |
| *C. tessellaris* | 7.04 ± 0.61 a | -2.39 | -1.93 | 43.3 | -4.48 ± 0.074 abc | 0.64 ± 0.13 de |
| *E. atrata* | 4.72 ± 0.88 ab | -1.61 | -1.36 | 24.67 | -5.54 ± 0.23 c | 2.9 ± 0.78 ab |
| *E. grandis* | 6.5 ± 0.89 ab | -2.11 | -1.8 | 31.51 | -4.67 ± 0.045 abc | 0.7 ± 0.20 cde |
| *E. laevopinea* | 5.95 ± 0.46 ab | -2.38 | -2.05 | 24.39 | -4.98 ± 0.96 bc | 1.75 ± 0.054 bcd |
| *E. longifolia* | 6.48 ± 0.67 ab | -2.36 | -2.3 | 25.22 | -3.82 ± 0.55 ab | 2.11 ± 0.29 ab |
| *E. macrandra* | 3.74 ± 0.15 b | -2.66 | -2.64 | 36.55 | -8.25 ± 0.51 d | 3.19 ± 0.49 a |
| *E. tereticornis* | 6.37 ± 0.94 ab | -2.63 | -2.4 | 43.41 | -3.5 ± 0.32 a | 0.49 ± 0.27 e |

### Relationships between hydraulic vulnerability, climate and SLA

Whilst diversity in in P50 and P12-P88 was observed across species, these traits appeared to be decoupled (linear relationship was not significant) (Figure 3 B). Leaf SLA and MAP had a weak linear relationship (P = 0.032, R^2^ = 0.56) (Figure 3 A). Significant linear relationships existed between P50 and leaf SLA (R^2^ = 0.75, **P < 0.01) but not with the elasticity of leaf tissues (ε) for the eight species of eucalypt (Figure 4). When drawing relationships between P50 and climate variables, the P50 was associated linearly with the MAP (mean annual precipitation) of a species’ home climate (R^2^ = 0.64, *P = 0.018) but was unrelated to the MAT (mean annual temperature) (Figure 4). P12-P88 also had a strong linear relationship with SLA (R^2^ = 0.72, **P <0.01), but unlike P50, the linear relationship with MAP was not significant (P > 0.05) (Figure 5). In context and generally speaking, leaves of species with more dry mass per unit area and from drier climates of origin reached P50 at more negative Ψ_leaf_ and spread cavitation over a greater range of water potentials than thinner leaves from species originating in wetter climates (Figure 4). Despite *E. laevopinea* leaf tissues being almost twice as hydraulically elastic than *C. tessellaris* and from a climate 10 degrees cooler on average, the P50 of the two species were comparable.

**Figure 3.**
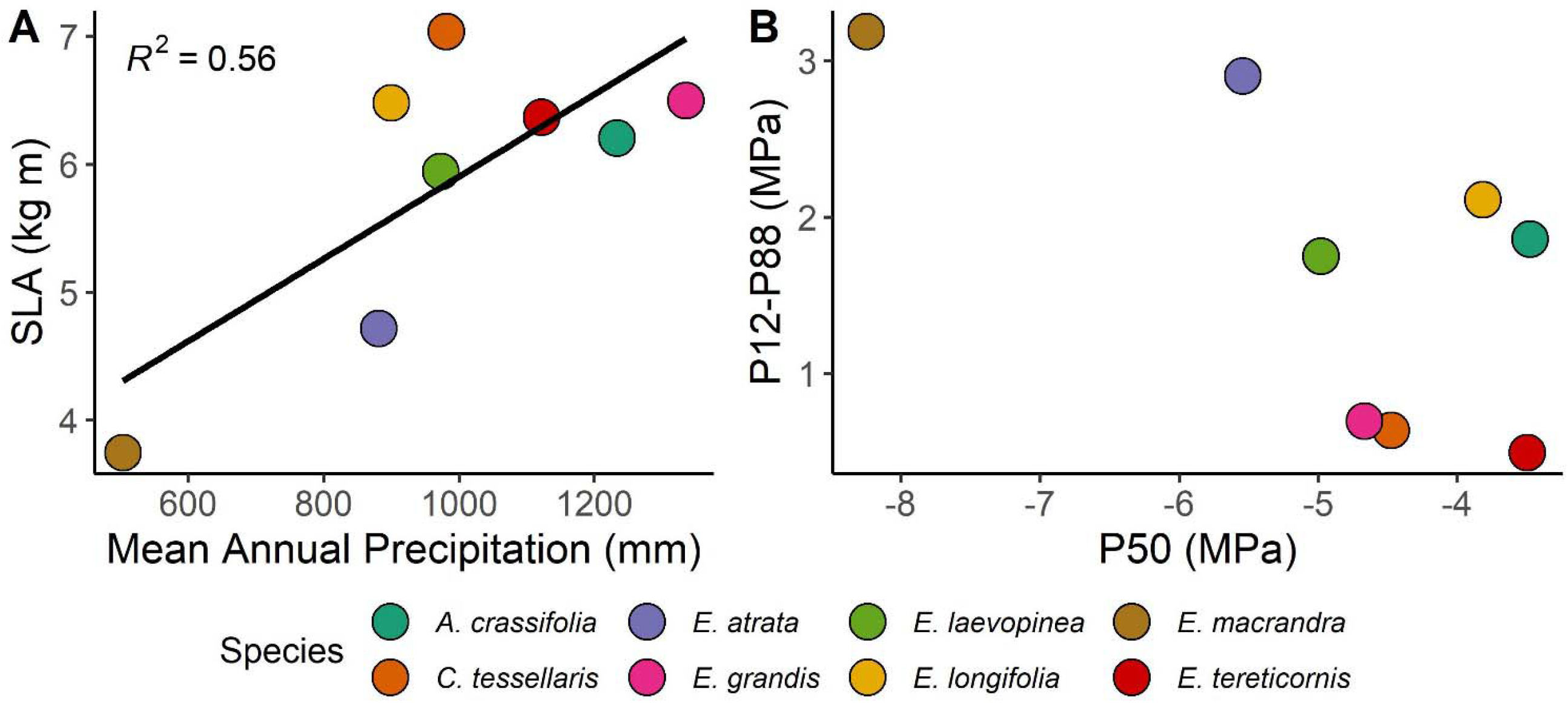
Relationships of leaf SLA and Mean Annual Precipitation (A) and two metrics of cavitation P50 and P12-P88 (B) of eight species of eucalypt. n = 4 for A. crassifolia and C. tessellaris, n = 3 for other species.

**Figure 4.**
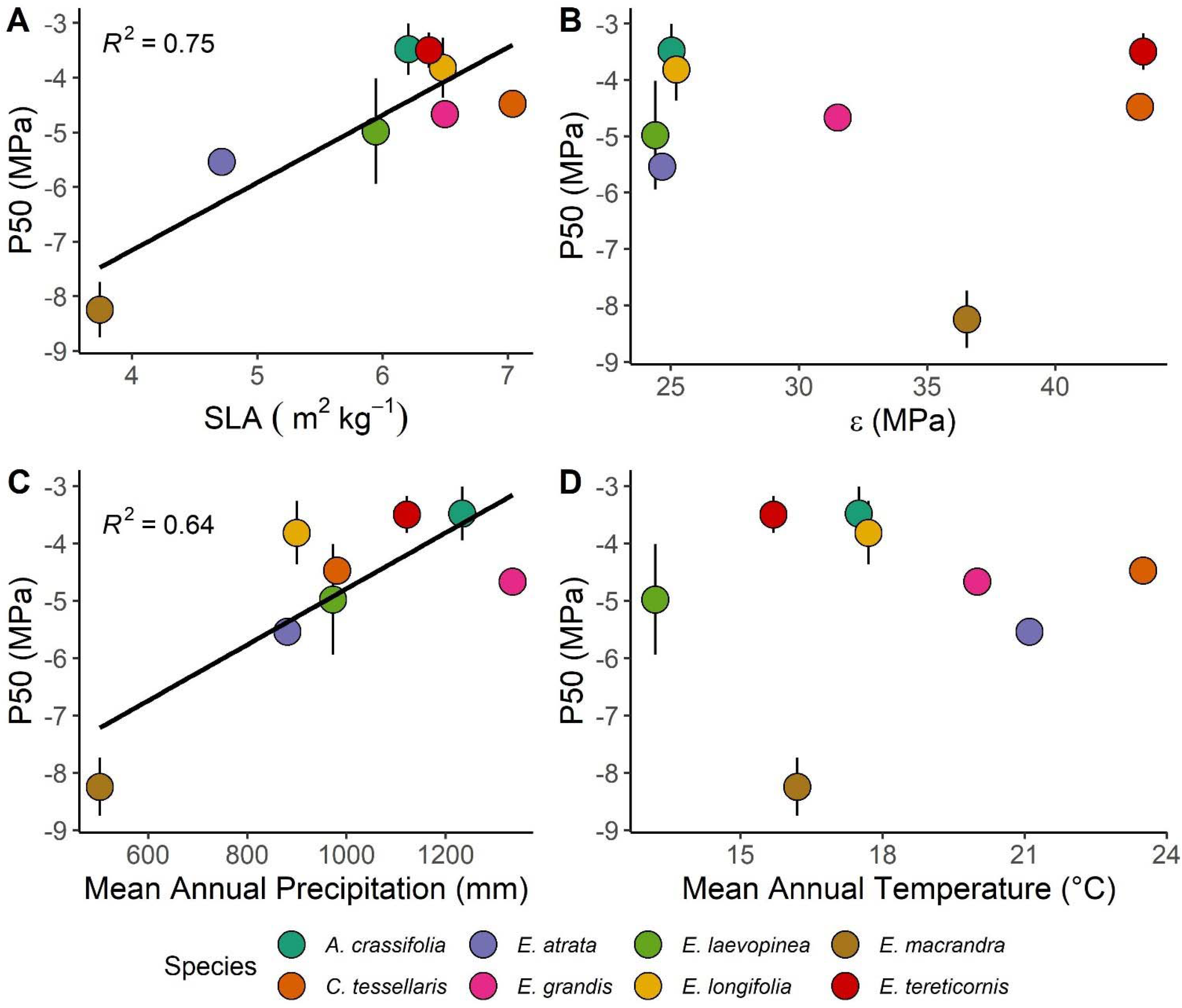
Relationships of P50 with leaf phsyiological variables of SLA (A), Bulk elastic modulus (B) and home climatic variables of precipitation (C) and temperature (D) of eight species of eucalypt. n = 4 for A. crassifolia and C. tessellaris, n = 3 for other species.

**Figure 5.**
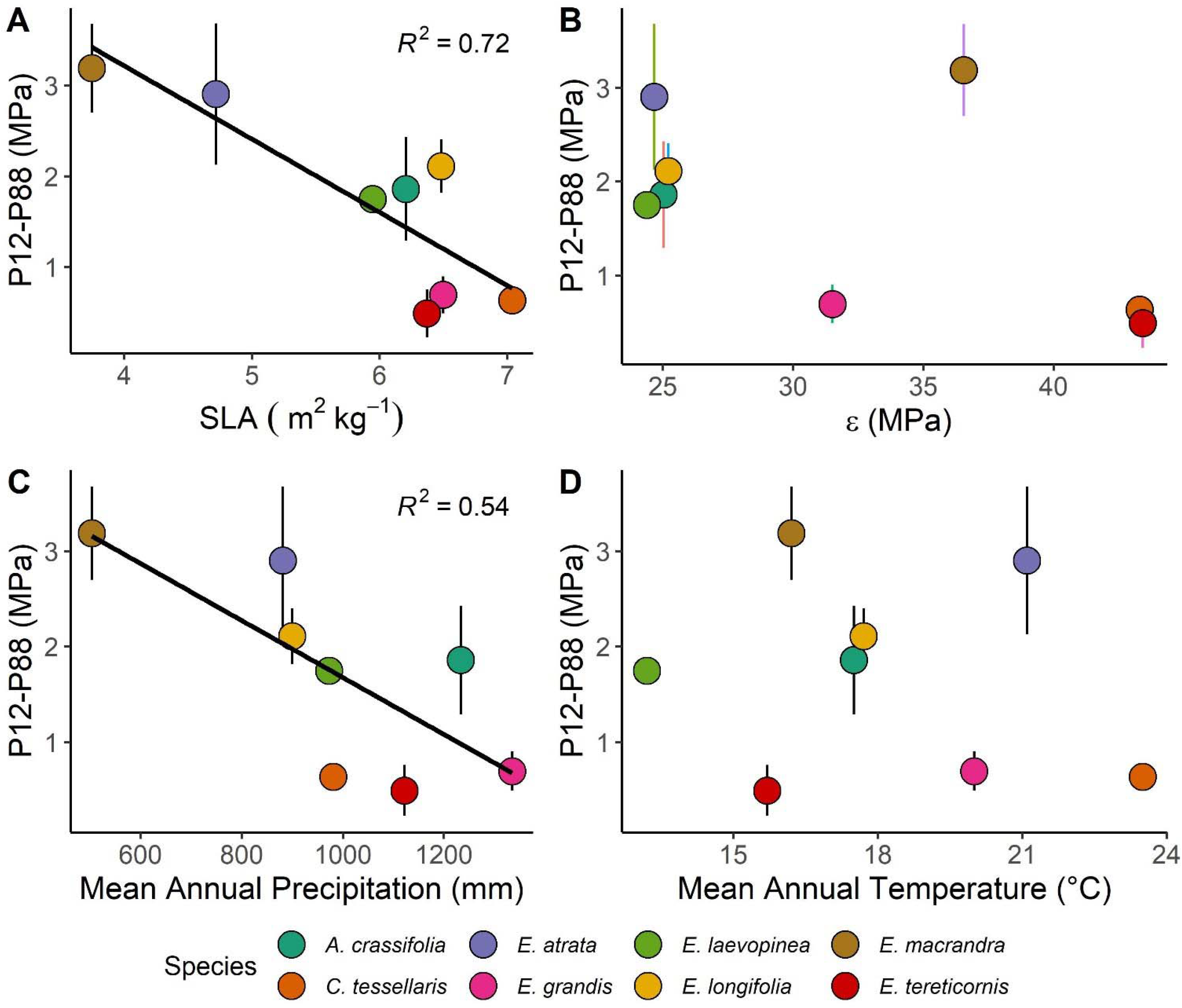
Relationships of P12-P88 with leaf phsyiological variables of SLA (A), Bulk elastic modulus (B) and home climatic variables of precipitation (C) and temperature (D) of eight species of eucalypt. n = 4 for A. crassifolia and C. tessellaris, n = 3 for other species

## Discussion

In our study, we measured the timing and spatial spread of cavitation through the leaf vein networks of eight eucalypt species and analysed how cavitation metrics correlated with climate of origin, leaf physiological and anatomical traits. Metrics of leaf hydraulic failure such as P50 and P12-P88 varied across species and were linked to specific leaf area and the annual precipitation at the climate of origin.

### Leaf cavitation progressed from larger to smaller conduits

Our results on the magnitude, as well as the sequence, of cavitation events in leaf tissue of eucalypts agreed well with other recent studies on similar species (Li *et al*., 2019; Creek *et al*., 2019). Cavitation began at less negative water potentials in the midveins and progressed to smaller, tertiary veins at increasingly negative water potentials (Li *et al*., 2019; Scoffoni *et al*., 2017b; Brodribb *et al*., 2016a). Although it could be argued that more distal veins would experience more negative water potentials earlier in the dry-down process and cavitate first, the vulnerability to cavitation of larger diameter vessels causes failure to occur first in situations where the water supply to the leaf is suddenly interrupted. The vein networks in leaves from all eucalypt species demonstrated a hierarchical, reticulate network with multiple, redundant conduits for the distribution of water across the mesophyll (Brodribb *et al*., 2016b). Under natural conditions, leaves of evergreen trees, including the eucalypt species in this study, experience cycles of hydraulic decline and recovery and must be able to maintain and recover mesophyll hydraulic conductivity. Redundancy in the major vein conduits allows water to circumvent embolisms which may preserve mesophyll tissue for longer and add resilience to the leaf hydraulic system. Interestingly, *E. macrandra* did not demonstrate cavitation of midvein conduits yet had the most embolism resistant leaves. It is possible that cavitation did occur here but was not visualised, due in part to poor transmission of light through the thickest part of these leaves – *E. macrandra* possessed the highest SLA of all the species studied. Certainly, visible cavitation in the major veins across the rest of the mesophyll in *E. macrandra* leaves preceded smaller veins, as was the case for other species. A commonly cited criticism of the OV method, although cheap, easy and non-destructive, is that not all cavitation events may be visible in leaves of all species due to poor resolution, superimposition of veins and imperfect transmission of light (Scoffoni *et al*., 2017b). Another limitation is that incorrect implementation of the method, particularly by incomplete or interrupted image capture, can produce vulnerability curves that are artificially vulnerable (Gauthey *et al*., 2020). The sparse appearance of cavitation across leaf vein networks for two species (*E. longifolia* and *E. laevopinea*) compared with the highly dense networks visible in *C. tessellaris*, may signify cavitation events in some of the smaller veins may not have been captured. However, *Eucalyptus* species from cooler, wetter climates do tend to have a much lower vein density (Brooker and Nicolle, 2013; de Boer *et al*., 2016), relying more heavily on outside xylem processes for water transport across the mesophyll. Furthermore, factors that control vein reticulation patterns in angiosperm leaf venation have been shown to be, in general, phylogenetically conserved at the genus level (Li *et al*., 2017). It may be that *C. tessellaris*, the most phylogenetically distinct species in this study (Thornhill *et al*., 2019), has a progression of cavitation that presents differently using the OV method.

### P50 and P12-P88 metrics were decoupled among species

In our study, P50 and P12-P88 were well-differentiated between species (Table 2), yet surprisingly, the rankings of the two parameters for these species did not correlate; the species with the lowest P50 did not spread the cavitation events over the largest range of water potentials (Figure 3 B). For example, *A. crassifolia* was first to lose half the visible conducting area, yet the cavitation process lasted longer than four other species. The role of P50 as an indicator of vulnerability to cavitation is well established, whereby leaves of species adapted to more arid conditions lie at the far end of the hydraulic safety/efficiency trade-off. These leaves have smaller conduit diameters, thicker cell walls, greater redundancy and low water transport resistance (Trueba *et al*., 2019; Grossiord *et al*., 2020). However, the range over which cavitation occurs, although intuitively a drought resistance trait, may relate more to the drought adaptive strategy of species, for example, where P50_stem_ is combined with leaf stomatal conductance traits to characterise iso/anisohydric strategies (Skelton *et al*., 2015). The hydraulic segmentation hypothesis applies to species with leaves that are drought avoiders – these species shed leaves during drought to reduce the evaporative demand and preserve conduit integrity in the stem (Pritzkow *et al*., 2019; Zimmermann, 2013; Pivovaroff *et al*., 2014a). The vein networks in these species may exhibit behaviour like those in *C. tessellaris* or *E. grandis* (Figure 2), where no damage is visible up to a certain point after which there is a rapid collapse of the system. In contrast, drought tolerators (Delzon, 2015) are species that retain their leaves and require the preservation of some conduits over a greater range of water potentials. Mesophyll tissue can continue to function, albeit with reduced efficiency, when water supply is restored to a partially cavitated leaf (Brodribb *et al*., 2016a). We hypothesise that the difference in strategy may be imparted by more (drought avoiders) or less (drought tolerators) uniform xylem conduit diameter sizes (Sack *et al*., 2015) across the network, causing air seeding in vessels to occur over a narrow range of water potentials. This is somewhat visible in our images of cavitation progression (Figure 1, vessels seem to be roughly uniform in size in *C. tessellaris* but have a range of sizes in *E. atrata*) although was not quantified in this study.

### The hydraulic margin between TLP and P50 highlights plastic versus genetically fixed hydraulic leaf traits in eucalypts

Osmotic adjustment of leaf tissue in response to water deficits is well described in a range of species, including eucalypts (Blum, 2017; Bartlett *et al*., 2012). In *Eucalyptus* species, this trait is remarkably flexible and can vary intra-specifically in response to local climates and temporally across seasons (Merchant *et al*., 2007). Leaves with higher concentrations of solutes in the vacuole can maintain leaf turgor despite greater strain on the hydraulic system, resulting in a shift of the TLP to lower water potentials. Our data agrees well with other common garden experiments in that TLPs are distributed over a relatively narrow range of water potentials (^~^ −1.5 to −3.5 MPa) and may reflect the convergence of osmotic adjustment among species as leaves optimise to the hydraulic conditions of the common climate (Warren *et al*., 2006; Li *et al*., 2019; Bourne *et al*., 2017; Li *et al*., 2018). In contrast, P50 was distributed over a much greater hydraulic range, suggesting this trait is more genetically fixed via permanent xylem structures during leaf development (Bourne *et al*., 2017). The need for a fixed expression of P50 is evidence that for leaves, there are serious consequences for plant growth following a loss of hydraulic integrity (Skelton *et al*., 2017).

A diverse range of Australian species, including eucalypts, appear to maintain stomatal conductance at water statuses below TLP; thus, TLP may not be as critical a threshold as P50 for these species (Farrell *et al*., 2017). Furthermore, outside xylem, hydraulic conductance rather than leaf xylem embolism has been shown to be the dominant cause of hydraulic decline at water potentials below TLP (Scoffoni *et al*., 2017c). A large hydraulic margin between TLP and P50, such as in *E. macrandra*, suggests that these species are evolutionarily adapted to dry climates but possess a TLP matched to relatively mild growing conditions.

### Cavitation metrics corresponded to precipitation of home climate and leaf SLA

Both P50 and P12-P88 correlated linearly with the mean annual precipitation (MAP) in the climate of seed origin. Many variations of these and other hydraulic traits characterising plant vulnerability to embolism have been demonstrated before in a range of species, including eucalypts. (Sack *et al*., 2015; Santiago *et al*., 2018; Pivovaroff *et al*., 2014a; Powell *et al*., 2017; Blackman *et al*., 2012; Blackman *et al*., 2009; Sack and Scoffoni, 2013; Bourne *et al*., 2017; Li *et al*., 2018; Liu *et al*., 2021). A mechanistic understanding of how species adapt to wetter climates by investing less in cavitation resistant leaf vein networks is well described by the efficiency-safety hypothesis (Santiago *et al*., 2018; Pivovaroff *et al*., 2014b); therefore, species adapted to lower rainfall possessing more negative P50 values is to be expected. The fact that P12-P88 did not correlate significantly with MAP supports our conclusion that this trait relates to drought strategy. The effect of decreasing MAP increasing the range of water potentials over which cavitation occurs gives rise to contrasting drought strategies in drought-avoiding leaves versus drought-tolerant leaves and highlights the critical role of resilience imparted by constitutive properties under strong genetic control. For *E. atrata*, cavitation events extended over three times as large a margin of water potentials than *C. tessellaris*, despite similar home climate MAPs. Neither P50 nor P12-P88 seemed to be affected by home climate MAT.

Evidence to suggest that leaf SLA may impart greater resistance to leaf vein cavitation among these eucalypt species was also observed (Figures 4 and 5). Low SLAs could increase resistance to cavitation in a number of ways, including more cellular layers which hold store water to support and buffer vein structures or greater investment in the xylem itself, including thicker xylem walls of vessels and greater vessel numbers per unit leaf area (Jordan *et al*., 2013; Sack *et al*., 2015). In contrast to our results, SLA did not present an advantage to P50 across several tropical tree species (Markesteijn *et al*., 2011). However, the eucalypts in this study had leaves with SLAs above these tropical species, exhibiting physical properties outside the scope of previous investigations. Furthermore, our data suggests that only SLAs below ^~^6 m^2^ kg^-1^ appear to confer a substantial advantage to extending P50 to lower water potentials among eucalypts. The similarity of SLA among species in this study (Table 2) may be the result of the interaction between the relatively mesic common garden environment and long-term adaption to a species’ home climate. Eucalypt leaves are typically highly plastic in form, and SLA may be somewhat plastic to suit the current growing environment in the six most mesic species (Brooker and Nicolle, 2013). The leaf morphology of *E. macrandra* and *E. atrata* may be limited in the extent to which investment in hydraulic architecture could match this environment, as these species’ home climates lay farthest geographically from the growing environment. SLA and investment in safe hydraulic structures may be a more fixed trait for these species. The significant correlation between longer P12-P88 and lower SLA points to investment in additional dry matter associated with hydraulic architecture (xylem or surrounding cellular structures) as causing more gradual rates of cavitation with declining hydraulic status (Li *et al*., 2017).

## Conclusion

Our results demonstrate how a detailed analysis of the hydraulic vulnerability of leaf vein networks among related species of eucalypts from contrasting climates can provide new insights into how and when cavitation resistance occurs and which metrics are coupled to home climate versus the growing climate. While P50 is correlated with home climate precipitation, the P12-P88 metric seems more related to drought strategy; more gradual cavitation progression with declining water content is represented by thicker leaves. Furthermore, our results agree with other studies that highlight P50 as a genetically fixed trait, while TLP may be more plastic, as demonstrated by similar TLP across these species grown in a common garden environment. Combining the OV method and vulnerability curves with other technologies, such as micro CT, have yielded a more detailed understanding of the still unresolved details of cavitation progression and recovery (Gauthey *et al*., 2020; Scoffoni *et al*., 2017b). Although we demonstrated a relationship with SLA and metrics of cavitation resistance, a better understanding of the underlying leaf anatomical structures between species that modulate P50 and P12-P88 could be achieved in future work using microscopy (Scoffoni *et al*., 2017b; Li *et al*., 2017). Similarly, applying the OV technique to a greater range of eucalypt species may further validate the trends shown in this study, particularly more species native to areas outside the high rainfall environments of south-eastern Australia.

---

**Supp. Figure 1.**
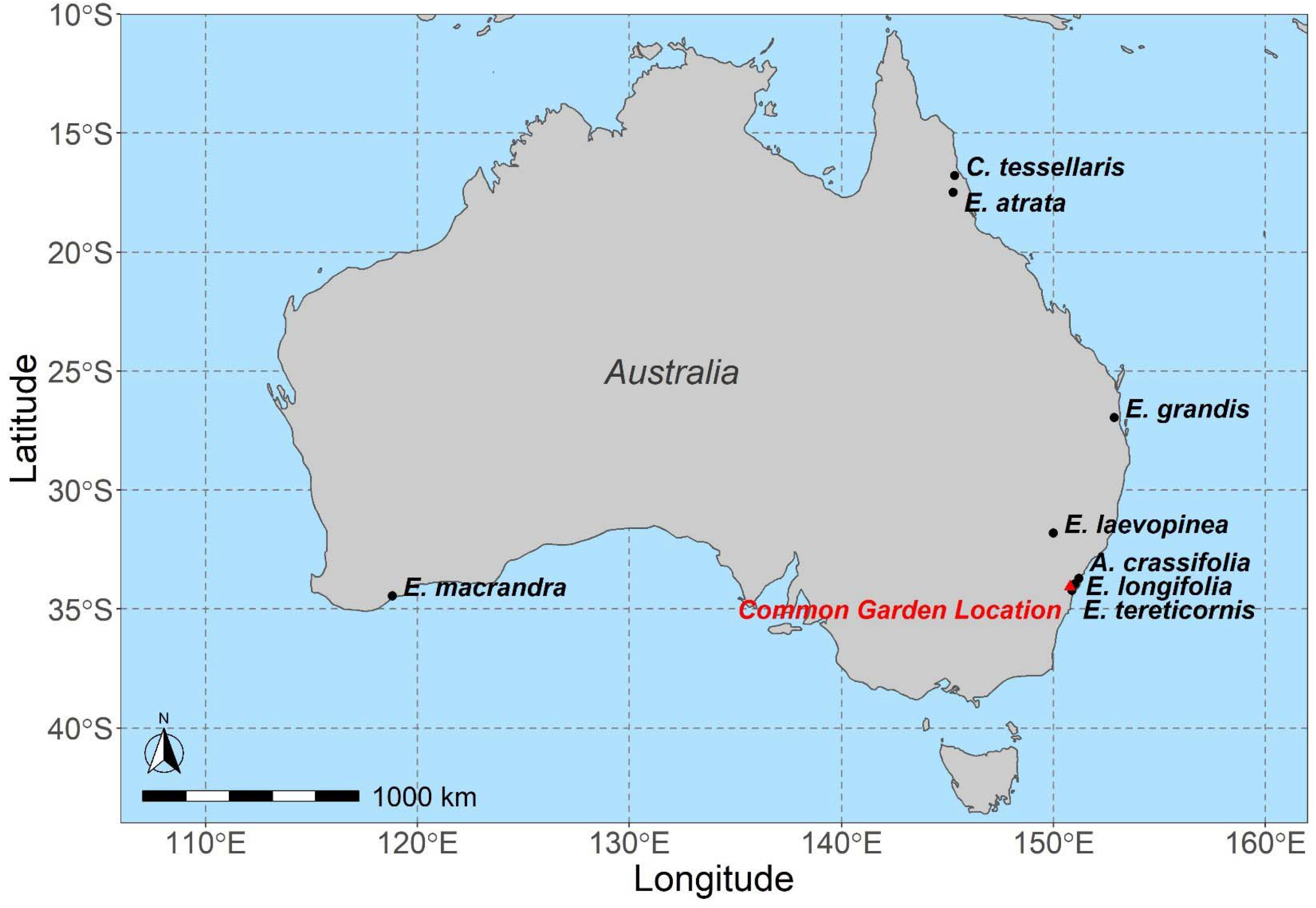
Map of seed collection locations for eight species of Eucalyptus from across Australia (See Table 1 for coordinates).

## Notes

### Competing Interest Statement

The authors have declared no competing interest.

## References

Aphalo, P. J. (2016). ggpmisc: An R package.

Bartlett, M. K., Scoffoni, C. & Sack, L. (2012). The determinants of leaf turgor loss point and prediction of drought tolerance of species and biomes: a global meta-analysis. Ecology Letters 15(5): 393–405.

Blackman, C. J., Aspinwall, M. J., Tissue, D. T. & Rymer, P. D. (2017). Genetic adaptation and phenotypic plasticity contribute to greater leaf hydraulic tolerance in response to drought in warmer climates. Tree Physiology 37(5): 583–592.

Blackman, C. J., Brodribb, T. J. & Jordan, G. J. (2009). Leaf hydraulics and drought stress: response, recovery and survivorship in four woody temperate plant species. Plant, Cell & Environment 32(11): 1584–1595.

Blackman, C. J., Brodribb, T. J. & Jordan, G. J. (2012). Leaf hydraulic vulnerability influences species’ bioclimatic limits in a diverse group of woody angiosperms. Oecologia 168(1): 1–10.

Blum, A. (2017). Osmotic adjustment is a prime drought stress adaptive engine in support of plant production. Plant, Cell & Environment 40(1): 4–10.

Bourne, A. E., Creek, D., Peters, J. M. R., Ellsworth, D. S. & Choat, B. (2017). Species climate range influences hydraulic and stomatal traits in Eucalyptus species. Annals of Botany 120(1): 123–133.

Brodribb, T. J., Bienaimé, D. & Marmottant, P. (2016a). Revealing catastrophic failure of leaf networks under stress. Proceedings of the National Academy of Sciences of the United States of America 113(17): 4865–4869.

Brodribb, T. J., Skelton, R. P., McAdam, S. A., Bienaimé, D., Lucani, C. J. & Marmottant, P. (2016b). Visual quantification of embolism reveals leaf vulnerability to hydraulic failure. New Phytologist 209(4): 1403–1409.

Brooker, I. & Nicolle, D. (2013). Atlas of leaf venation and oil gland patterns in the eucalypts. Csiro publishing.

Choat, B., Brodribb, T. J., Brodersen, C. R., Duursma, R. A., López, R. & Medlyn, B. E. (2018). Triggers of tree mortality under drought. Nature 558(7711): 531–539.

Creek, D., Lamarque, L. J., Torres-Ruiz, J. M., Parise, C., Burlett, R., Tissue, D. T. & Delzon, S. (2019). Xylem embolism in leaves does not occur with open stomata: evidence from direct observations using the optical visualization technique. Journal of Experimental Botany 71(3): 1151–1159.

de Boer, H. J., Drake, P. L., Wendt, E., Price, C. A., Schulze, E.-D., Turner, N. C., Nicolle, D. & Veneklaas, E. J. (2016). Apparent Overinvestment in Leaf Venation Relaxes Leaf Morphological Constraints on Photosynthesis in Arid Habitats Plant Physiology 172(4): 2286–2299.

de Mendiburu, F. & de Mendiburu, M. F. (2020). Package ‘agricolae’. R package version: 1.2-8.

Delzon, S. (2015). New insight into leaf drought tolerance. Functional Ecology 29(10): 1247–1249.

Dixon, M. & Downey, A. (2013). PSY1 Stem Psychrometer manual. ICT International [online].

Farrell, C., Szota, C. & Arndt, S. K. (2017). Does the turgor loss point characterize drought response in dryland plants? Plant, Cell & Environment 40(8): 1500–1511.

Gauthey, A., Peters, J. M. R., Carins-Murphy, M. R., Rodriguez-Dominguez, C. M., Li, X., Delzon, S., King, A., López, R., Medlyn, B. E., Tissue, D. T., Brodribb, T. J. & Choat, B. (2020). Visual and hydraulic techniques produce similar estimates of cavitation resistance in woody species. New Phytologist 228(3): 884–897.

Grossiord, C., Ulrich, D. E. & Vilagrosa, A. (2020). Controls of the hydraulic safety–efficiency trade-off. Tree Physiology 40(5): 573–576.

Hartmann, H., Moura, C. F., Anderegg, W. R. L., Ruehr, N. K., Salmon, Y., Allen, C. D., Arndt, S. K., Breshears, D. D., Davi, H., Galbraith, D., Ruthrof, K. X., Wunder, J., Adams, H. D., Bloemen, J., Cailleret, M., Cobb, R., Gessler, A., Grams, T. E. E., Jansen, S., Kautz, M., Lloret, F. & O’Brien, M. (2018). Research frontiers for improving our understanding of drought-induced tree and forest mortality. New Phytologist 218(1): 15–28.

Johnson, D. M., Berry, Z. C., Baker, K. V., Smith, D. D., McCulloh, K. A. & Domec, J.-C. (2018). Leaf hydraulic parameters are more plastic in species that experience a wider range of leaf water potentials. Functional Ecology 32(4): 894–903.

Johnson, D. M., Meinzer, F. C., Woodruff, D. R. & Mcculloh, K. A. (2009). Leaf xylem embolism, detected acoustically and by cryo-SEM, corresponds to decreases in leaf hydraulic conductance in four evergreen species. Plant, Cell & Environment 32(7): 828–836.

Jordan, G. J., Brodribb, T. J., Blackman, C. J. & Weston, P. H. (2013). Climate drives vein anatomy in Proteaceae. American journal of botany 100(8): 1483–1493.

Kassambara, A. (2020). rstatix: Pipe-friendly framework for basic statistical tests. R package version 0.6. 0.

Klein, T., Zeppel, M., Anderegg, W., Bloemen, J., De Kauwe, M., Hudson, P., Ruehr, N., Powell, T., von Arx, G. & Nardini, A. (2018). Xylem embolism refilling and resilience against drought-induced mortality in woody plants: processes and trade-offs. Ecological Research.

Li, L., Ma, Z., Niinemets, Ü. & Guo, D. (2017). Three Key Sub-leaf Modules and the Diversity of Leaf Designs. Frontiers in Plant Science 8(1542).

Li, X., Blackman, C. J., Choat, B., Duursma, R. A., Rymer, P. D., Medlyn, B. E. & Tissue, D. T. (2018). Tree hydraulic traits are coordinated and strongly linked to climate-of-origin across a rainfall gradient. Plant, Cell & Environment 41(3): 646–660.

Li, X., Blackman, C. J., Peters, J. M. R., Choat, B., Rymer, P. D., Medlyn, B. E. & Tissue, D. T. (2019). More than iso/anisohydry: Hydroscapes integrate plant water use and drought tolerance traits in 10 eucalypt species from contrasting climates. Functional Ecology 33(6): 1035–1049.

Liu, H., Ye, Q., Gleason, S. M., He, P. & Yin, D. (2021). Weak tradeoff between xylem hydraulic efficiency and safety: climatic seasonality matters. New Phytologist 229(3): 1440–1452.

Markesteijn, L., Poorter, L., Paz, H., Sack, L. & Bongers, F. (2011). Ecological differentiation in xylem cavitation resistance is associated with stem and leaf structural traits. Plant, Cell & Environment 34(1): 137–148.

Merchant, A., Callister, A., Arndt, S., Tausz, M. & Adams, M. (2007). Contrasting physiological responses of six Eucalyptus species to water deficit. Annals of Botany 100(7): 1507–1515.

Pau, G., Fuchs, F., Sklyar, O., Boutros, M. & Huber, W. (2010). EBImage—an R package for image processing with applications to cellular phenotypes. Bioinformatics 26(7): 979–981.

Pivovaroff, A. L., Sack, L. & Santiago, L. S. (2014a). Coordination of stem and leaf hydraulic conductance in southern California shrubs: a test of the hydraulic segmentation hypothesis. New Phytologist 203(3): 842–850.

Pivovaroff, A. L., Sack, L. & Santiago, L. S. (2014b). Coordination of stem and leaf hydraulic conductance in southern California shrubs: a test of the hydraulic segmentation hypothesis. New Phytologist 203(3): 842–850.

Powell, T. L., Wheeler, J. K., de Oliveira, A. A. R., da Costa, A. C. L., Saleska, S. R., Meir, P. & Moorcroft, P. R. (2017). Differences in xylem and leaf hydraulic traits explain differences in drought tolerance among mature Amazon rainforest trees. Global Change Biology 23(10): 4280–4293.

Pritzkow, C., Williamson, V., Szota, C., Trouvé, R. & Arndt, S. K. (2019). Phenotypic plasticity and genetic adaptation of functional traits influences intra-specific variation in hydraulic efficiency and safety. Tree Physiology 40(2): 215–229.

R Core Team (2019). R: A language and environment for statistical computing. In R Foundation for Statistical Computing, Vol. 2020 Vienna, Austria.

Rodriguez-Dominguez, C. M., Carins Murphy, M. R., Lucani, C. & Brodribb, T. J. (2018). Mapping xylem failure in disparate organs of whole plants reveals extreme resistance in olive roots. New Phytologist 218(3): 1025–1035.

Rueden, C. T., Schindelin, J., Hiner, M. C., DeZonia, B. E., Walter, A. E., Arena, E. T. & Eliceiri, K. W. (2017). ImageJ2: ImageJ for the next generation of scientific image data. BMC bioinformatics 18(1): 529.

Sack, L. & Holbrook, N. M. (2006). Leaf Hydraulics. Annual review of plant biology 57(1): 361–381.

Sack, L. & Scoffoni, C. (2013). Leaf venation: structure, function, development, evolution, ecology and applications in the past, present and future. New Phytologist 198(4): 983–1000.

Sack, L., Scoffoni, C., Johnson, D. M., Buckley, T. N. & Brodribb, T. J. (2015). The Anatomical Determinants of Leaf Hydraulic Function. In Functional and Ecological Xylem Anatomy, 255–271 (Ed U. Hacke). Cham: Springer International Publishing.

Santiago, L. S., De Guzman, M. E., Baraloto, C., Vogenberg, J. E., Brodie, M., Hérault, B., Fortunel, C. & Bonal, D. (2018). Coordination and trade-offs among hydraulic safety, efficiency and drought avoidance traits in Amazonian rainforest canopy tree species. New Phytologist 218(3): 1015–1024.

Schindelin, J., Arganda-Carreras, I., Frise, E., Kaynig, V., Longair, M., Pietzsch, T., Preibisch, S., Rueden, C., Saalfeld, S. & Schmid, B. (2012). Fiji: an open-source platform for biological-image analysis. Nature methods 9(7): 676–682.

Scoffoni, C., Albuquerque, C., Brodersen, C. R., Townes, S. V., John, G. P., Bartlett, M. K., Buckley, T. N., McElrone, A. J. & Sack, L. (2017a). Outside-Xylem Vulnerability, Not Xylem Embolism, Controls Leaf Hydraulic Decline during Dehydration. Plant Physiology 173(2): 1197–1210.

Scoffoni, C., Albuquerque, C., Brodersen, C. R., Townes, S. V., John, G. P., Cochard, H., Buckley, T. N., McElrone, A. J. & Sack, L. (2017b). Leaf vein xylem conduit diameter influences susceptibility to embolism and hydraulic decline. New Phytologist 213(3): 1076–1092.

Scoffoni, C., Albuquerque, C., Cochard, H., Buckley, T. N., Fletcher, L. R., Caringella, M. A., Bartlett, M., Brodersen, C. R., Jansen, S., McElrone, A. J. & Sack, L. (2018). The Causes of Leaf Hydraulic Vulnerability and Its Influence on Gas Exchange in Arabidopsis thaliana. Plant Physiology 178(4): 1584.

Scoffoni, C., Sack, L. & Ort, D. (2017c). The causes and consequences of leaf hydraulic decline with dehydration. Journal of Experimental Botany 68(16): 4479–4496.

Skelton, R. P., Brodribb, T. J., McAdam, S. A. M. & Mitchell, P. J. (2017). Gas exchange recovery following natural drought is rapid unless limited by loss of leaf hydraulic conductance: evidence from an evergreen woodland. New Phytologist 215(4): 1399–1412.

Skelton, R. P., West, A. G. & Dawson, T. E. (2015). Predicting plant vulnerability to drought in biodiverse regions using functional traits. Proceedings of the National Academy of Sciences 112(18): 5744–5749.

Team RStudio (2015). RStudio: integrated development for R. RStudio Inc., Boston, MA 42: 14.

Thornhill, A. H., Crisp, M. D., Külheim, C., Lam, K. E., Nelson, L. A., Yeates, D. K. & Miller, J. T. (2019). A dated molecular perspective of eucalypt taxonomy, evolution and diversification. Australian Systematic Botany 32(1): 29–48.

Trueba, S., Pan, R., Scoffoni, C., John, G. P., Davis, S. D. & Sack, L. (2019). Thresholds for leaf damage due to dehydration: declines of hydraulic function, stomatal conductance and cellular integrity precede those for photochemistry. New Phytologist 223(1): 134–149.

Venturas, M. D., Sperry, J. S. & Hacke, U. G. (2017). Plant xylem hydraulics: what we understand, current research, and future challenges. Journal of Integrative Plant Biology 59(6): 356–389.

Villanueva, R. A. M. & Chen, Z. J. (2019). ggplot2: Elegant graphics for data analysis. Taylor & Francis.

Warren, C., Dreyer, E., Tausz, M. & Adams, M. (2006). Ecotype adaptation and acclimation of leaf traits to rainfall in 29 species of 16-year-old Eucalyptus at two common gardens. Functional Ecology 20(6): 929–940.

Zhu, S. D., Liu, H., Xu, Q. Y., Cao, K. F. & Ye, Q. (2016). Are leaves more vulnerable to cavitation than branches? Functional Ecology 30(11): 1740–1744.

Zimmermann, M. H. (2013). Xylem structure and the ascent of sap. Springer Science & Business Media.

